# Mechanical Properties of DNA Double-Crossover Motif

**DOI:** 10.1101/2023.02.28.530469

**Authors:** Eva Matoušková, Michal Růžička, Kamila Réblová, Filip Lankaš

**Affiliations:** Department of Informatics and Chemistry, University of Chemistry and Technology Prague, Czech Republic; CEITEC – Central European Institute of Technology, Brno, Czech Republic

## Abstract

The double-crossover (DX) motif is a key building block of DNA nanostructures. It connects two double helices by two closely spaced Holliday junctions. Despite its prominent importance, the structure and elasticity of the DX motif is not fully understood. Here we employ extensive all-atom molecular dynamics (MD) simulations of an antiparallel DNA DX motif with two full turns between crossovers to infer its global structure and deformability. We quantitatively reproduce the experimentally known two-fold increase of bending stiffness upon incorporation of a DNA duplex into the DX motif, and find out that its stretching and twisting stiffness are only slightly influenced. To further describe the motif, we define four effective rigid bodies, each involving several base pairs flanking the Holliday junctions from the outside, and consider internal coordinates capturing relative displacement and orientation of the bodies. Time series of the coordinates from MD then yield the global structure of the DX motif and its stiffness in the harmonic approximation.

---

DNA double-crossover molecules, investigated 30 years ago (1), have become a prominent building block of DNA nanostructures (2). They comprise two DNA double helices linked together by two strand exchange points. There are several types of DX motifs, differing in the relative orientation of the two duplexes, which may be either parallel or antiparallel, and by the even or odd number of half-turns between the crossovers. The most promising type for nanotechnology application is the DAE motif, comprising antiparallel helices with an even number of half-turns between the crossovers. This type, specifically the one containing two turns of DNA between its crossover points, has been investigated by a ligation cyclization assay (3), and it was found that the bending stiffness of a DNA duplex increases roughly two-fold if a second duplex is fused to it to form a DX motif. However, the effect on other duplex stiffness characteristics remains largely unknown. Little is also known about the shape and stiffness of the motif as a whole, not just of one duplex.

In this work we set out to investigate global mechanical properties of a DAE motif by means of unrestrained all-atom MD simulations. Our system includes a DX motif comprising two 46 base-pair (bp) DNA duplexes connected by two crossing points 20 bp apart (Fig. 1A), surrounded by explicitly represented water molecules and ions in an octahedral periodic box. K+ ions added to neutralize the DNA charge are supplemented by additional K+ and Cl-ions to mimic the 150 mM monovalent salt concentration. The OL15 force field for DNA (4), Dang parameters for the ions (5) and the SPC/E water model are used. After a series of energy minimizations and short MD runs to equilibrate the system, a 1 µs trajectory has been produced. The 4 bp portions of the duplexes flanking the crossing points from outside are each assigned an effective rigid body, as described (6). Briefly, each of the 8 bases involved in a body are fitted with a standard reference point and frame (7). The rigid body reference point is then obtained as the arithmetic mean of the base reference points, while the rigid body frame is computed as the Euclidean average of the base frames (8) and further rotated so that its z-axis coincides with the mean helical axis of the fragment. An isolated DNA duplex of the same sequence has been also simulated as a control.

**Figure 1.**
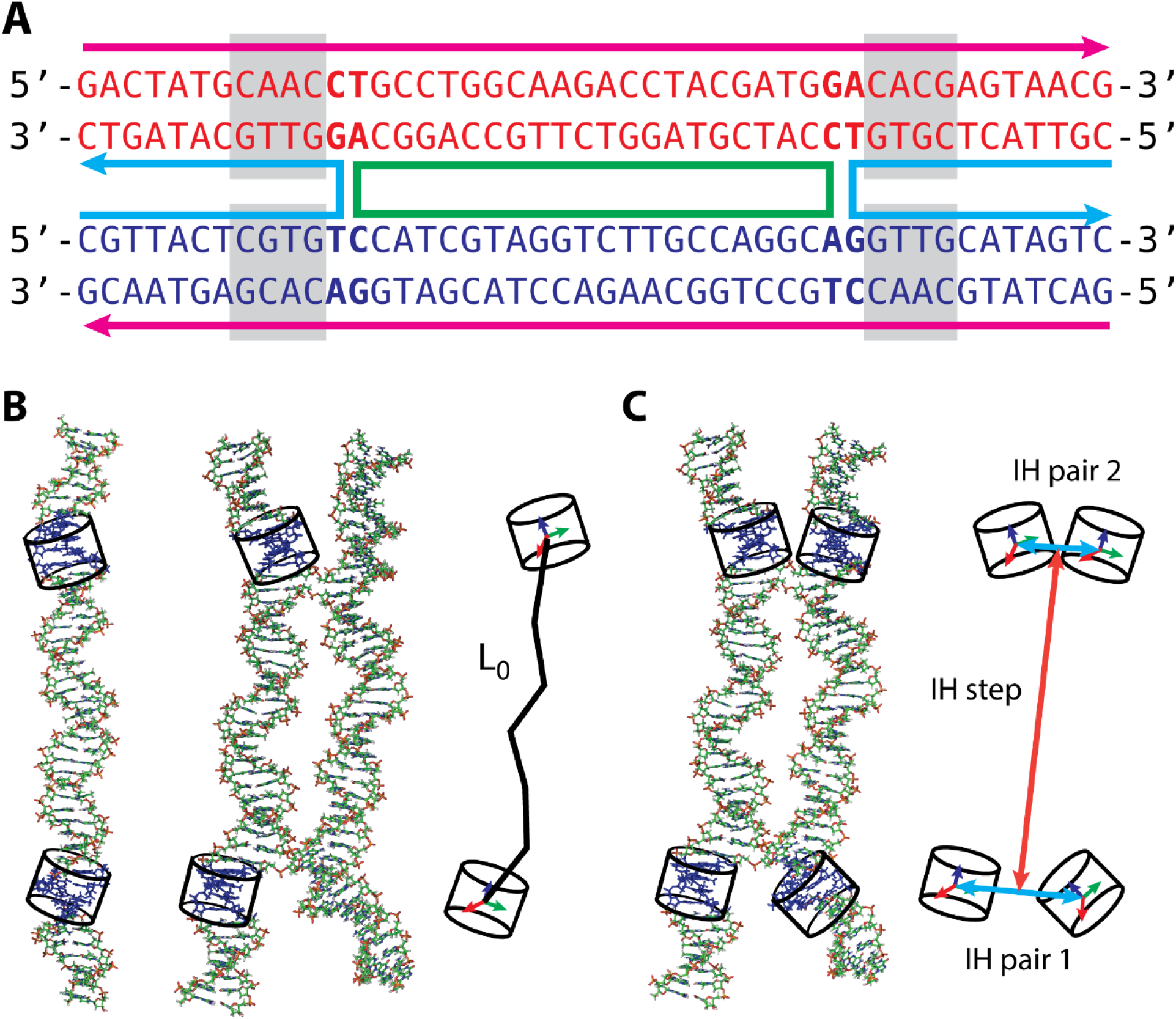
(A) Sequence and schematic representation of the DX motif studied in this work. (B) An MD snapshot of the reference duplex and of the motif with the effective rigid bodies, obtained as averages of the reference points and frames of four bases flanking the crossover points. The length and the total twist is measured between the bodies. (C) A global description of the motif using four rigid bodies whose relative displacement and rotation are defined using 18 internal coordinates, six for each inter-helical (IH) pair and six for the IH step. The coordinate definitions are analogous to the common local intra-base pair and step coordinates used to capture the conformation of a DNA duplex.

We first focus on just one duplex (Fig. 1B). To compute its stretch modulus, we define the duplex length as a sum of the helical rises between the reference points of the two bodies (half the value of the helical rise is taken in the innermost base-pair step within each body). Assuming a quadratic dependence of the deformation energy on elongation (Hooke’s law), the stretch modulus *Y* is related to the variance *σ*^2^ of the length by the usual relation (see for example here (9))

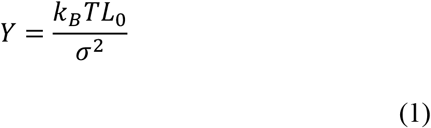

where *L*_0_is the equilibrium length computed as the mean length over the MD trajectory, *k*_*B*_ is the Boltzmann constant and *T* the absolute temperature at which the simulations were performed (300 K).

Similarly, substituting the variance of the total twist, computed as a sum of helical twists between the reference points of the two bodies (once again, half the value is taken in the innermost steps), into eq. 1 yields the twist rigidity. It is convenient to divide it by *k*_*B*_*T* to obtain its value in the units of length. We did the computation for the free duplex and for each of the two duplexes in the DX motif. Since all three of them have an identical base sequence, they are directly comparable. Moreover, the two duplexes in the motif should behave in an identical manner for symmetry reasons.

To assess the bending rigidity of the duplex, we define a frame in the middle between the two end frames and compute the bending angles into the x- and y-directions of this middle frame, designated as the global roll (Ro) and global tilt (Ti) respectively. The definition is exactly analogous to the one of the local roll and tilt as employed by the 3DNA conformational analysis algorithm (10). In this case we have a 2-dimensional problem, i.e. two coordinates, and we assume once again the elastic energy being a quadratic form characterized by the equilibrium values of the coordinates and by the stiffness matrix *K* which, in turn, is computed from the coordinate covariance matrix *V* using the relation (9)

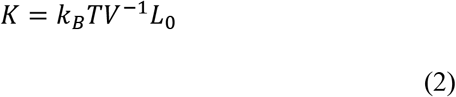

It is again convenient to divide the matrix elements of *K* by *k*_*B*_*T* to express them in the units of length. Besides the diagonal stiffness constants *A*_*Ro*_ and *A*_*Ti*_, we also have the coupling term *A*_*RT*_.

The results are in Table 1. The very small value of the coupling term *A*_*RT*_ enables us to characterize the bending stiffness by the quantity *A*_*iso*_, the effective bending rigidity of a hypothetical isotropic elastic rod, defined by the relation (11)

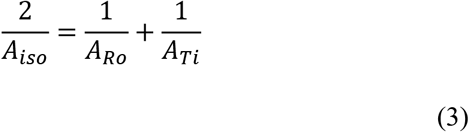

**Table 1.** Elastic constants of a free duplex and of the two duplexes within the DX motif.

| | $A_{Ro}$ (nm) | $A_{Ti}$ (nm) | $A_{RT}$ (nm) | $A_{iso}$ (nm) | $Y$ (pN) | $C$ (nm) |
| --- | --- | --- | --- | --- | --- | --- |
| Duplex | $67 \pm 2$ | $74 \pm 3$ | $0 \pm 5$ | $70 \pm 0$ | $1602 \pm 29$ | $131 \pm 6$ |
| Dx1 | $126 \pm 6$ | $231 \pm 6$ | $-34 \pm 11$ | $163 \pm 6$ | $1407 \pm 28$ | $146 \pm 16$ |
| Dx2 | $114 \pm 4$ | $219 \pm 8$ | $-38 \pm 11$ | $150 \pm 1$ | $1453 \pm 28$ | $150 \pm 3$ |

The bending stiffness of the free duplex agrees quantitatively with the DNA dynamic persistence length of 78 ± 13 nm found in cryo-electron microscopy experiments (12), the stretch modulus and twist rigidity are slightly higher than the experimental values of 1000-1500 pN and 100 nm, respectively (13). However, here we are more interested in the differential effect due to incorporating the duplex into the DX motif. The first thing to note is that the two duplexes in the motif behave quite similarly, an important check of the convergence of our MD. Comparing their elastic moduli with those of the free duplex, we see that the built-in duplexes are indeed slightly more than twice as stiff in bending, exactly as found in the experiment (3). Moreover, our data indicate that the DX duplexes are also somewhat more flexible in stretching and more rigid in twisting, but these changes are by far not as dramatic as the effect on bending.

We further investigate the global conformation and stiffness of the DX motif. To this end we define four rigid bodies, each of them at the outer part of the duplex just flanking the crossing point (Fig. 1C). To define their relative displacement and rotation, we proceed in close analogy with the usual description of base pairs and steps in a DNA double helix (10). Thus, we have two inter-helical (IH) pairs, effective “base pairs”, each characterized by the six standard intra-base pair coordinates buckle, propeller, opening, shear, stretch and stagger, and one IH step described by the six standard step coordinates tilt, roll, twist, shift, slide and rise. We use the 3DNA definitions (10) throughout, with one important exception. Each duplex is itself twisted, so that the orientation of the rigid body frames would change rapidly with the position of the bodies along the duplex. To avoid this, we define the “base-pair” frames not just by averaging the two rigid body frames as done in 3DNA, but using a construction summarized in Fig. 3. Briefly, we project the vector connecting the two reference points onto the plane perpendicular to the mean z-vector, then define the y-axis of the middle frame as the unit vector along this projection. Thus, the connecting vector now lies in the yz-plane of the frame and, consequently, shear (i.e. its projection onto the x-axis) is exactly zero. In this way we eliminate the effect of rapid rotation of the body frames as the bodies are shifted along the duplexes, at the expense of loosing one degree of freedom per IH pair.

**Figure 2.**
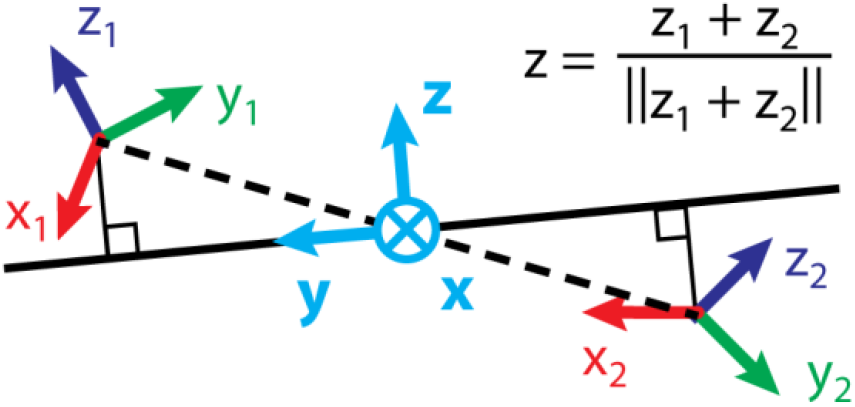
Construction of the reference frame in the middle between the two rigid bodies constituting the inter-helical (IH) pair, an effective “base pair”.

**Figure 3.**
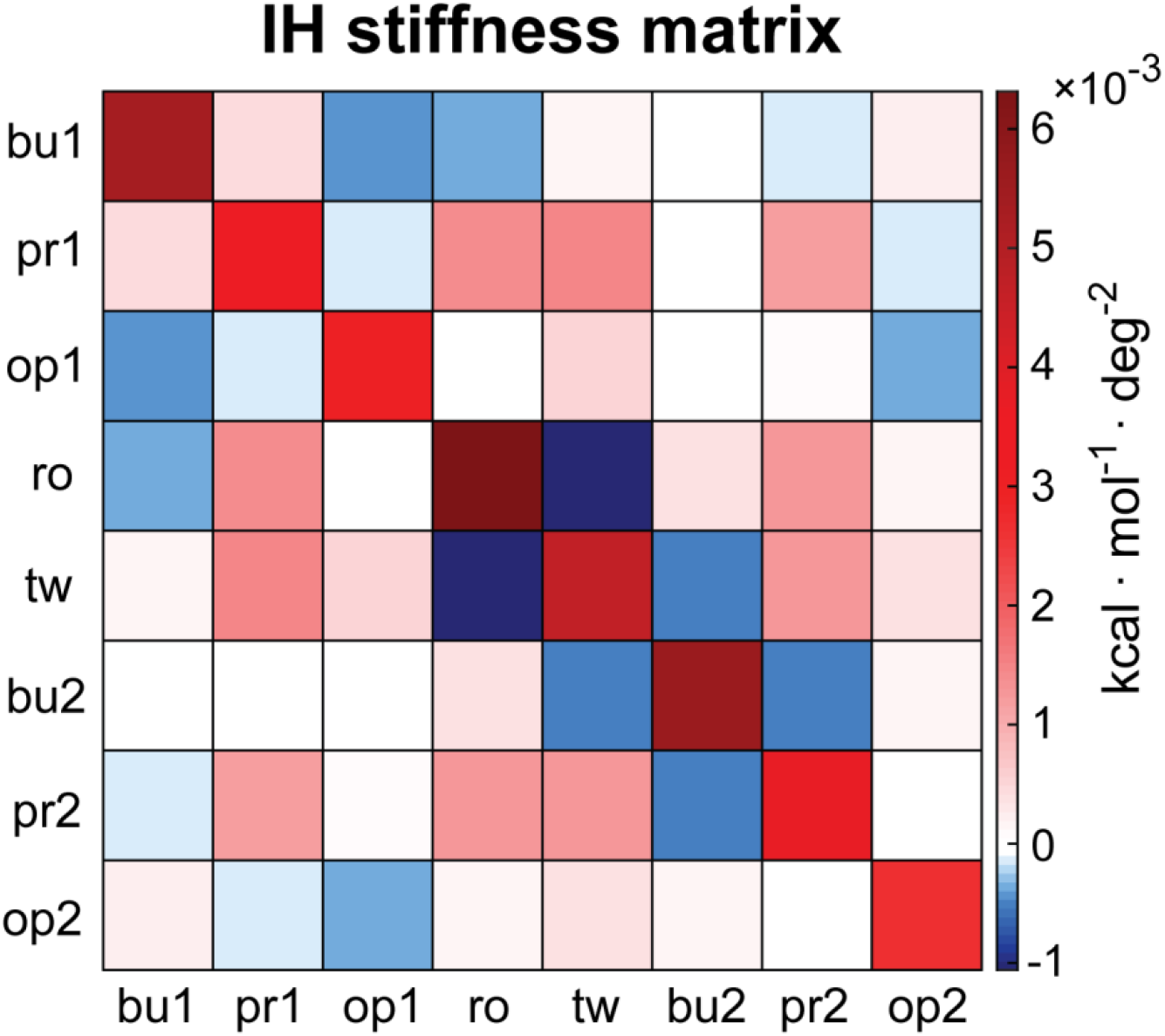
Stiffness matrix for selected inter-helical (IH) coordinates. All the other IH coordinates are much stiffer than these and are excluded from the calculation.

We again assume the deformation energy to be a quadratic form of these IH coordinates (there are 16 of them altogether), containing the equilibrium values of the coordinates and the stiffness matrix as its parameters. The latter is computed from the IH coordinate covariance as

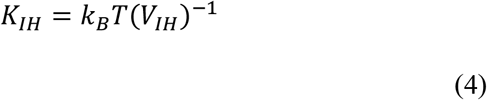

Notice that here we do not normalize the stiffness according to the length of the structure: the stiffness constants are therefore valid for the given DX motif and may change if, for instance, the number of helical turns between the crossing points changes.

The average coordinates are in Table S1. We observe that the DX motif is slightly bent (by ca. 10°) and twisted (∼37°), while the two duplexes are not exactly parallel – rather, they are slightly rotated about the x-axis of the IH pair frame (Fig. 3) by 17° (IH buckle) and about its y-axis (IH propeller) and, more importantly, the duplexes are rotated about the z-axis as well, so that their grooves do not point into the same direction but point ca. 58° away (IH opening).

As for the stiffness, to compare translational and angular degrees of freedom, we have to make them dimensionally uniform. To do so, we observe that a rotation by *φ* radians about the IH pair z-axis will displace the rigid body reference points by *rφ*/2, where *r* is the distance between the duplexes. Since we observe *r* around 25 Å, this gives a relation between the length and the angle scale and enables us to compare the stiffness of the different degrees of freedom directly. The comparison shows that all of the translational degrees of freedom and the tilt (i.e. bending towards the duplexes) and far stiffer than the other coordinates. They are therefore omitted from the analysis. The remaining ones include the three rotational degrees of freedom of each IH pair plus the IH roll (i.e. bending perpendicular to the plane of the motif) and IH twist. Thus, we are left with 8 coordinates altogether. The stiffness matrix is in Fig. 3, the values are listed in Table S2. The values would enable one to compute the energetic cost of any deformation of the DX motif at this level of description.

Notice finally that the particular choice of the ordering of the IH pairs is arbitrary. If we followed the motif in the opposite direction, then the IH pairs would be interchanged and all the frames rotated by 180° about their x-axes. Upon this transformation, the “odd” IH coordinates (buckle, shear, tilt, shift) change sign, while the remaining “even” ones remain unchanged. Consequently, the odd means (Table S1) and the stiffness constants (Table S2) corresponding to an odd-even combination change sign, while all the other parameters stay the same. Thus, the coordinate choice made here enables one to easily switch between the two descriptions.

In summary, we propose a method to characterize global mechanical properties of a DNA DX motif in terms of relative displacements and rotations of its flanking stems. A short part of each stem adjacent to the crossing point is modelled as an effective rigid body, and internal coordinates are defined to capture the relative spatial arrangement of the bodies. We assume a quadratic deformation energy function and deduce its parameters, namely the equilibrium geometry and the stiffness matrix of the motif, from all-atom MD simulations. Our results quantitatively reproduce the experimentally known stiffening of a DNA duplex with respect to bending upon its incorporation into the motif, and further indicate that, in contrast to bending, the duplex stretching and twisting stiffness are only slightly affected. Moreover, we provide a full characterization of the DX motif shape and stiffness in terms of the spatial configuration of its flanking duplexes. Our results may contribute to better understanding of the shape and stiffness properties of this key nanostructure building block.

## Acknowledgment

This work was supported from the grant of Specific University Research – grant no. A2_FCHT_046 to EM and FL.

## Supplementary Tables

**Table S1.**
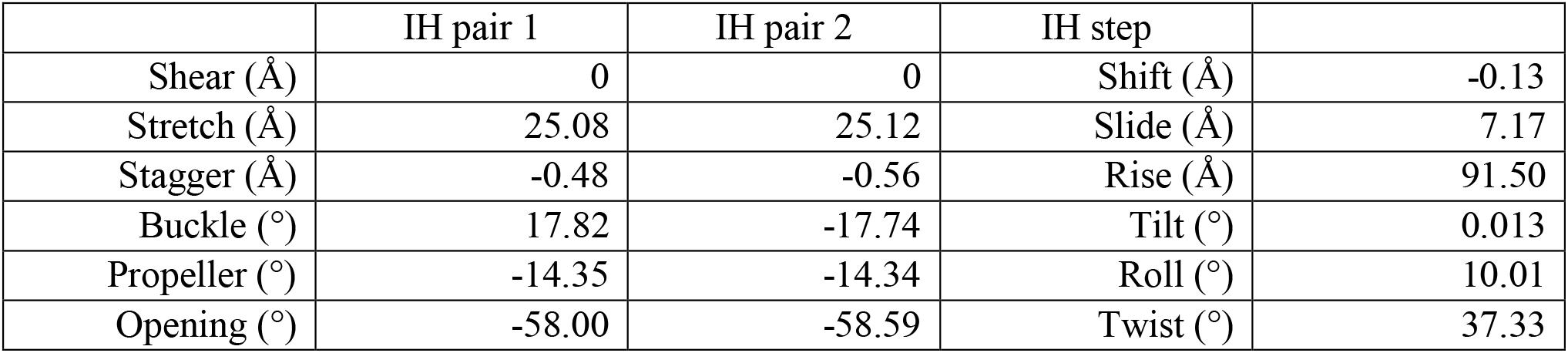
Equilibrium values of the IH coordinates for the DX motif.

|  | IH pair 1 | IH pair 2 | IH step |  |
| --- | --- | --- | --- | --- |
| Shear (Å) | 0 | 0 | Shift (Å) | -0.13 |
| Stretch (Å) | 25.08 | 25.12 | Slide (Å) | 7.17 |
| Stagger (Å) | -0.48 | -0.56 | Rise (Å) | 91.50 |
| Buckle (°) | 17.82 | -17.74 | Tilt (°) | 0.013 |
| Propeller (°) | -14.35 | -14.34 | Roll (°) | 10.01 |
| Opening (°) | -58.00 | -58.59 | Twist (°) | 37.33 |

**Table S2.** Elements of the DX motif IH stiffness matrix.

|  |  |  |  |  |  |
| --- | --- | --- | --- | --- | --- |
| **bu1-bu1** | 5.35 | **op1-pro2** | 0.086 | **bu2-tw** | -0.53 |
| **bu1-pro1** | 0.43 | **op1-op2** | -0.34 | **bu2-bu2** | 5.59 |
| **bu1-op1** | -0.42 | **ro-bu1** | -0.38 | **bu2-pro2** | -0.48 |
| **bu1-ro** | -0.38 | **ro-pro1** | 1.43 | **bu2-op2** | 0.13 |
| **bu1-tw** | 0.19 | **ro-op1** | 0.02 | **pro2-bu1** | -0.14 |
| **bu1-bu2** | -0.08 | **ro-ro** | 6.32 | **pro2-pro1** | 1.16 |
| **bu1-pro2** | -0.14 | **ro-tw** | -1.06 | **pro2-op1** | 0.09 |
| **bu1-op2** | 0.24 | **ro-bu2** | 0.34 | **pro2-ro** | 1.28 |
| **pro1-bu1** | 0.43 | **ro-pro2** | 1.28 | **pro2-tw** | 1.28 |
| **pro1-pro1** | 3.57 | **ro-op2** | 0.16 | **pro2-bu2** | -0.48 |
| **pro1-op1** | -0.11 | **tw-bu1** | 0.19 | **pro2-pro2** | 3.68 |
| **pro1-ro** | 1.43 | **tw-pro1** | 1.47 | **pro2-op2** | -0.06 |
| **pro1-tw** | 1.47 | **tw-op1** | 0.49 | **op2-bu1** | 0.24 |
| **pro1-bu2** | 0.04 | **tw-ro** | -1.06 | **op2-pro1** | -0.11 |
| **pro1-pro2** | 1.16 | **tw-tw** | 4.69 | **op2-op1** | -0.34 |
| **pro1-op2** | -0.11 | **tw-bu2** | -0.53 | **op2-ro** | 0.16 |
| **op1-bu1** | -0.42 | **tw-pro2** | 1.28 | **op2-tw** | 0.38 |
| **op1-pro1** | -0.11 | **tw-op2** | 0.38 | **op2-bu2** | 0.13 |
| **op1-op1** | 3.02 | **bu2-bu1** | -0.08 | **op2-pro2** | -0.06 |
| **op1-ro** | 0.02 | **bu2-pro1** | 0.04 | **op2-op2** | 2.63 |
| **op1-tw** | 0.49 | **bu2-op1** | -0.02 |  | x 10<sup>-3</sup><br>kcal/mol/deg<sup>2</sup> |
| **op1-bu2** | -0.02 | **bu2-ro** | 0.34 |  |  |

